# Network-based molecular subtyping of acral melanoma

**DOI:** 10.1101/2023.02.04.527155

**Authors:** Yin Mingzhu, Yiding Zhang, Wenhua Wang, Shuang Zhao, Juan Su, Shao Li, Xiang Chen

## Abstract

Acral melanoma is more biologically aggressive with a worse prognosis compared with other melanoma subtypes. However, the molecular basis underlying the biological and clinical behavior of this cancer is still unclear. Here, using the combination of multi-omics data analysis and network-based disease gene prediction algorithm, we first demonstrate the existence of two acral melanoma subtypes which greatly differed in clinical performance, cellular and molecular mechanisms, and discovered a biomarker panel (EREG, VSIG4, FCGR3A, RAB20) that accurately distinguished these two subtypes with the AUC of 0.946, which has been verified by clinical samples. Subtype I has thinner Breslow with a better prognosis. On the contrary, subtype II is a high-risk subtype that is easier to invade the dermis. We further analyzes the intrinsic biological mechanism of the two subtypes from the cellular level, and reveals the important role of macrophages subgroups in the molecular typing of acral melanoma. Feature genes of subtype I are enriched in FCN1+ macrophages that promote inflammatory and immune responses. In contrast, feature genes of subtype II are enriched in SPP1+ macrophages which ha the greatest impact on tumor cells. The identification of the two subtypes opens up important biological and clinical perspectives for acral melanoma.

## 1 Introduction

Traditional tumor typing is generally based on the location, histopathology, and clinical features of diseases. However, tumors are also highly heterogeneous in molecular biology [1] and the application of tumor molecular typing is of great significance for accurate disease diagnosis and drug discovery [2–4]. Acral melanoma occurs on hairless skin that is not easily exposed to sunlight, such as the palms, soles, and under the nails [5]. Acral melanoma has many unique features that distinguish it from other types of cutaneous melanoma(CM), including delayed discovery, high malignancy, and the greater tendency to metastasize after malignant transformation, which contribute to a poorer associated prognosis and survival rate than other CM subtypes [6, 7]. Due to the heterogeneity, treatment and prognostic outcomes may also vary widely among patients with acral melanoma[5, 8]. Therefore, it is necessary to discover disease subtypes with distinct molecular mechanisms and identify driving factors and key gene pathways for individualized treatment.

Up to now, some works have been carried out on molecular typing of cutaneous melanoma. Liu et al[9] proposed a single-sample immune signature network and identified an immune-related subtype based on hierarchical clustering of transcriptome data. The immune-related subtype were further subdivided into three subtype, one of which have the highest degree of immune infiltration and generally have better outcomes after immunotherapy. Tsoi et al [10] used the data of human melanoma cell lines and discovered four cell groups which have large differences in the differentiation patterns. In addition, there is also related works on uveal melanoma[11, 12]. For example, Robertson et al [13] performed unsupervised clustering of 80 uveal melanomas in TCGA based on somatic copy number variation data and classified them into four subtypes with distinct molecular mechanisms and clinical prognosis.

Although works in cutaneous and uveal melanoma have demonstrated that studying the heterogeneity based on molecular features is critical for personalized treatment, fine-grained molecular typing of acral melanoma is still lacking. Moreover, most large-scale studies only use traditional bulk data for unsupervised clustering. Though machine learning based methods such as CIBERSORT[14], xCell[15], ESTIMATE can estimate the cell type proportion from bulk data, it is still no substitute for high-resolution interpretation of cellular information from single-cell sequencing data.

Therefore, a new computational framework for the molecular typing of acral melanoma is constructed (Fig. 1). The combination of bulk RNA-seq and single-cell sequencing data overcomes the problem of insufficient single-cell data sample size due to the high cost of sequencing, while taking into account the use of high-precision cell-level information, avoiding noises from sampling and traditional sequencing. In addition, to reduce the limitations brought by the heterogeneity of omics data and sample sizes, we introduce network-based causative gene prediction algorithms that provide a systematic and holistic perspective for understanding the underlying mechanisms of complex diseases. Finally, two novel subtypes of acral melanoma and their corresponding markers were discovered, which were validated by clinical samples. We verified the differences between the two subtypes from both clinical features and molecular mechanisms, and proved that subtype II is a high-risk subtype with a higher degree of malignancy. Based on single-cell data analysis, it was further revealed that subgroups of macrophages with different functions play a vital role in acral melanoma typing.

**Fig. 1.**
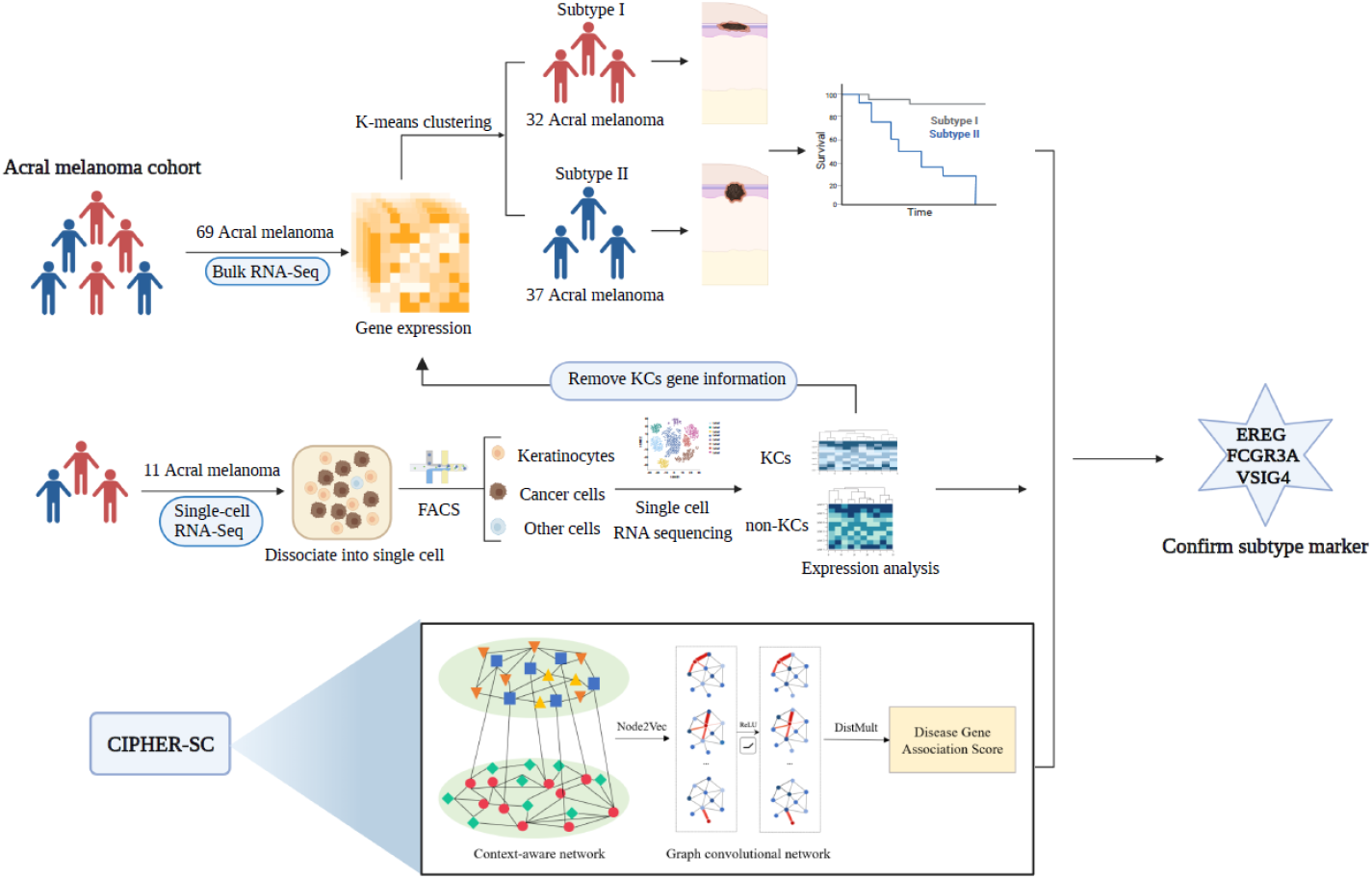
The process of molecular subtyping analysis. Acral melanoma is divided into two subtypes and the subtyping markers are identified through the combined analysis of bulk RNA-seq, single-cell RNA-seq, and network-based prediction algorithm.

## 2 Results

### 2.1 Identification of two subtypes of acral melanoma based on unsupervised clustering

To analyze the heterogeneity of acral melanoma, we collect 69 tumor samples from acral melanoma patients from multiple centers. We use the next-generation high-throughput NovaSeq 6000 RNA-Seq platform to generate RNA-seq data and the data after quality control is retained for subsequent analysis.

We first calculate the Euclidean distance between sample pairs to analyze the similarity between acral melanoma patients. It can be found that the transcriptome expression pattern of all samples is heterogeneous, suggesting that the molecular characteristics is important to identify new subtypes (Fig. 2a). We use K-Means to perform unsupervised clustering of samples based on transcriptome data. By setting the K value to 2, 3, 4 and 5 in turn to cluster the samples, we then use Principal Component Analysis (PCA) to reduce the dimensionality and visualize the clustering results (Fig. 2b). Gap statistic and Silhouette Coefficient are used to evaluate the clustering effect as K takes different values (Fig. 2c). The samples are finally categorized into two subtypes: subtype I contains 32 patients, and subtype II contains 37 patients.

**Fig. 2.**
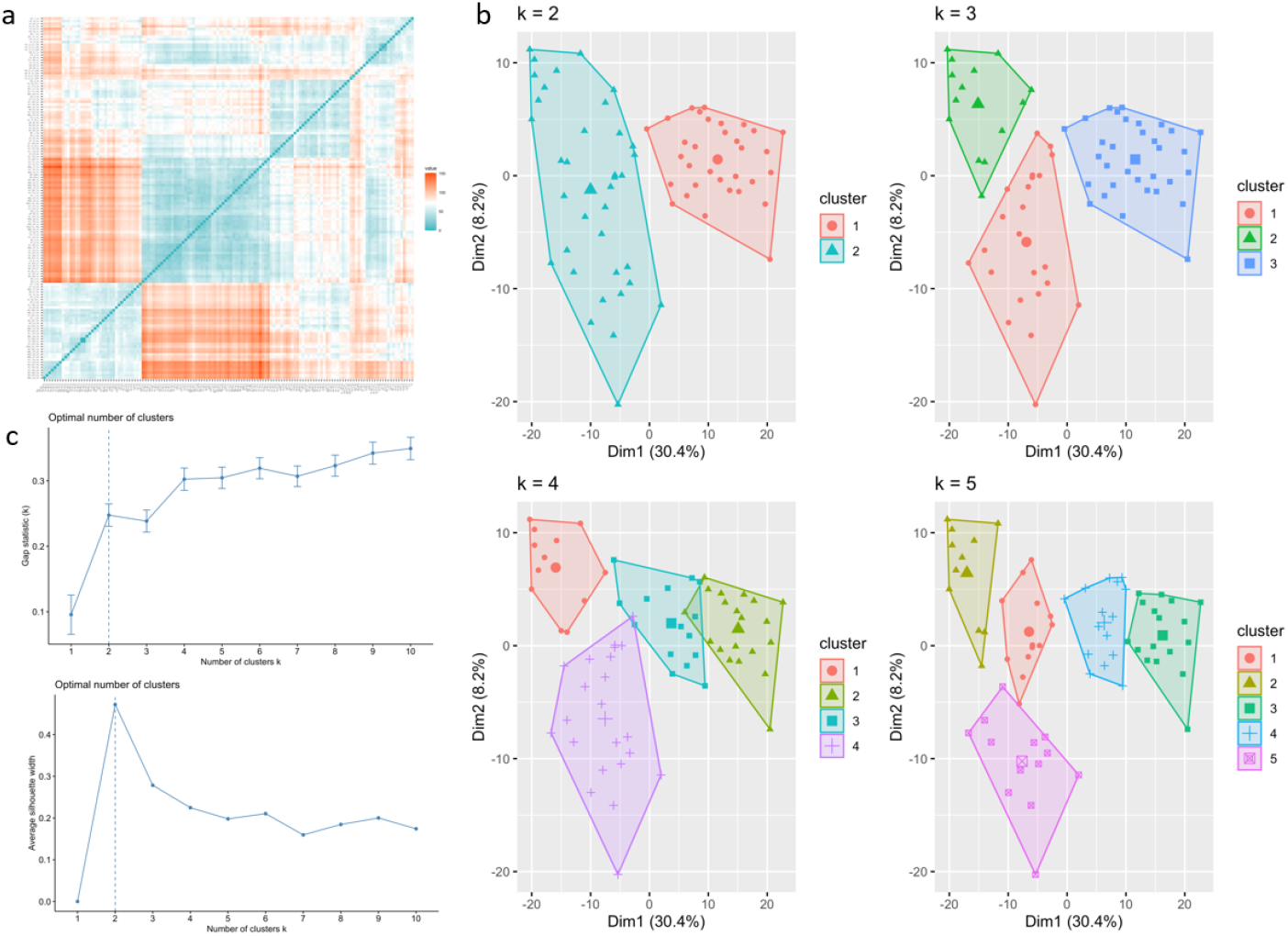
Heterogeneity and subtypes identification of acral melanoma. **a** The Euclidean distance between every pair of samples. Blue indicates nearer distance while orange means farther distance. **b** Unsupervised clustering results when K takes 2,3,4 and 5 using the K-means algorithm. K means the number of clusters.**c** Gap statistic and Silhouette width value at different number of clusters. The optimal number of clusters is marked with dotted line.

### 2.2 The subtypes are significantly different both from molecular and cellular level

To compare the differences between these two subtypes, differential expression analysis is then carried out, with 642 genes significantly up-regulated in subtype I and 300 genes in subtype II (*FDR* < 0.05, *log*_2_*FC* > 1) (Fig. 3a). The enriched GO functions are analyzed respectively. The enriched GOs in subtype I include skin development (*P* < 2.3*e* − 28), epidermal cell differentiation (*P* < 2.3*e* − 26), lipid catabolic process (*P* < 3.7*e* − 05), establishment of skin barrier (*P* < 8.2*e* − 05) and etc (Fig. 3c). Since melanoma is limited to the uppermost epidermis in the early stage, the thickness of the tumor will increase and gradually invade into the deeper dermis as the tumor develops. The results indicated that the main characteristics of subtype I were the dysfunction of normal epidermal cells, the destruction of the skin barrier, lipid metabolism and other related functions.

**Fig. 3.**
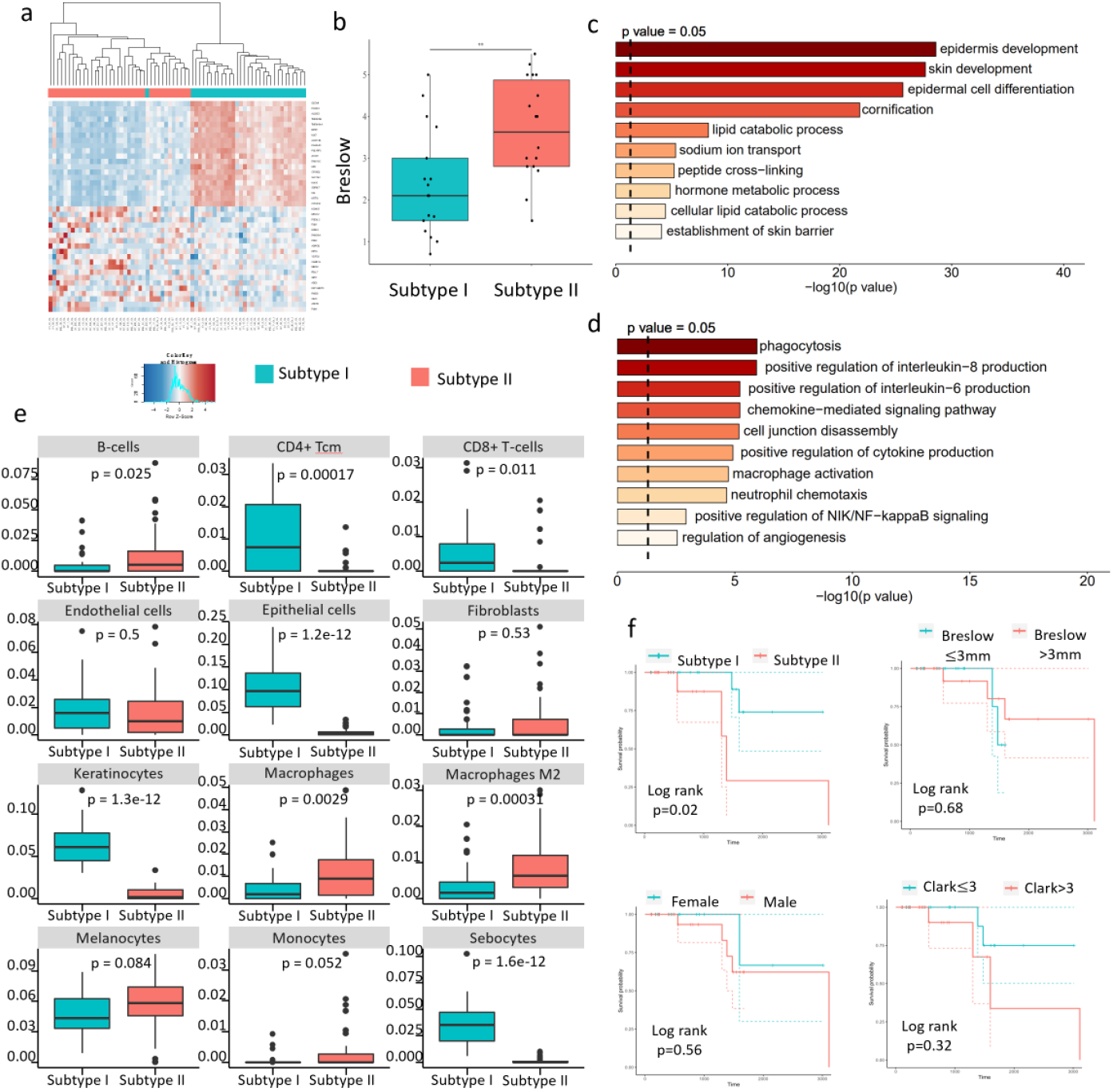
The two subtypes differ significantly at both the micro and macro levels. **a** Heatmap of differentially expressed genes between two subtypes of acral melanoma. **b** Breslow thickness boxplot of subtype I and subtype II. **c**,**d** The enriched GO functions in subtype I and subtype II respectively. **e** The enrichment scores of each cell type are calculated by Xcell and are then compared to discover the celllular composition difference betweeen two subtypes. **f** The prognostic effects of new subtypes and other clinical characteristics including breslow thickness, gender, and clark’s class.

The enriched GOs in subtype II include phagocytosis (*P* < 2.9 *e* −7), positive regulation of interleukin-8 production (*P* < 1.2*e* −6), chemokine-mediated signaling pathway (*P* < 6.1*e* −6), cell junction disassembly (*P* < 6.7*e* −6), macrophage activation (*P* < 1.9*e* −5), positive regulation of *NIK/NF − kappaB* signaling (*P* < 1.2*e* −3), regulation of angiogenesis (*P* < 2.9*e* −3) and etc (Fig. 3d). These functions are mainly related to cell connection changes, chemokine production, angiogenesis, and tumorigenesis. CXCL8 is an important immunosuppressive factor, which can be secreted by tumor cells, tumor-associated fibroblasts, macrophages and other cells[16].

Studies have shown that its expression level is closely related to the progression of melanoma[17], which is considered to be one of the most effective angiogenic factors.

From the above enrichment analysis results, it can be seen that the two subtypes may have large differences in the tumor microenvironment. Therefore, we further analyze the composition of cell types for the two subtypes. To resolve cell-level information, we use the Xcell to infer the enrichment scores of different types of cells for the two subtypes[15]. It can be found that the enrichment scores of epithelial cells, keratinocytes, CD4+ central memory T cells and CD8+ T cells in subtype I are significantly higher (Fig. 3e). Among them, the significant enrichment of keratinocytes may be due to the thinner tumor thickness in tumor samples of subtype I, and thus a higher proportion of keratinocytes in the uppermost layer of the skin are obtained at the time of sampling. On the contrary, subtype II is significantly enriched with immune cells such as macrophages and B cells, especially M2 type macrophages (*P* < 3*e* −4). It can be seen that immune cells enriched in subtype I are mainly related to tumor suppression, while immune cells in subtype II are related to tumorigenesis. This result suggests that the immune microenvironment play an important role in the classification of acral melanoma, and is worthy of further study.

In addition, TCM syndromes are often used in the classification and treatment of diseases[18, 19]. We find that the previously reported cold and hot-related marker genes(CSF1R, CXCL8)[20, 21] are related to the differentially expressed genes of these two subtypes, and the deeper association needs further research.

### 2.3 Subtype II is a high-risk subtype with a worse prognosis

We then compared the clinical characteristics of the two subtypes. Breslow thickness measures the distance from the top of the granular layer of the skin epidermis to the deepest invasive cells in the tumor. A large number of studies have shown that there is a direct relationship between Breslow thickness and patient survival, and it is the most important clinical prognostic feature of melanoma tumors. The American Joint Committee on Cancer (AJCC) proposed a cutoff of 1mm and pointed out that melanomas thinner than it usually have better prognosis, while melanomas above this threshold incline to have a worse prognosis [22, 23]. As depicted in Fig. 3b, patients in subtype II generally have significantly thicker Breslow thickness (*P* < 0.0059) and the median reaches 3.6mm. It indicates that tumor in subtype II has broken through the epidermis of the skin and invaded into the dermis and even the subcutaneous tissue.

We further compare the differences of survival between two subtypes, and analyze the effect of existing clinical characteristics on the prognosis of patients. KM survival curves show significant differences of the survival of the two subtypes, subtype II have a significantly worse prognosis (*P* < 0.02, *logranktest*). We also analyze the prognostic effects of other clinical characteristics such as Breslow thickness, gender, and Clark’s class (Fig. 3f). The results demonstrate that these clinical characteristics have a poor prognostic effect on acral melanomas. Although patients with higher Clark’s class have a worse trend on the KM survival curve, the difference is not statistically significant in the log-rank test. Above all, the new subtype has a vital significance for the clinical prognosis.

To evaluate the impacts of multiple risk factors on survival, cox proportional hazards regression analysis is further applied to deal with continuous values or multi-category variables. The hazard ratio of the new classification is greater than 1 with P smaller than 0.05 whether in univariate analysis or multivariate analysis, indicating that there is still a strong correlation between patients of subtype II and shorter survival time in the absence of other factors (Table. 1). In contrast, the impact of other variables is not significant.

### 2.4 Identification of subtype markers

We have proved that the subtypes are distinct in molecular functions and clinical characteristics, demonstrating that subtype I is a low-risk subtype confined to the epidermis and subytpe II is a high-risk subtype prone to invasion. To better distinguish the subytpes with different risks and achieve personalized treatments, we then identified biomarkers of these two subytpes based on the applicaiton of single-cell data and network-based prediction methods. The mechanisms at the molecular and cellular levels are further analyzed as well.

#### 2.4.1 A single-cell transcriptomic atlas of acral melanoma

In order to explore the heterogeneity and analyze marker functions at single-cell resolution, we constructed a single-cell atlas of acral melanoma based on 11 acral melanoma samples. BD Rhapsody platform is used to generate single-cell RNA-seq data. After quality control and data processing, 110647 cells are obtained for subsequent analysis. To identify distinct cell populations, dimension reduction and unsupervised clustering are performed using methods implemented in Seurat. Ten cell types are identified and later annotated by canonical markers (Fig. 4a,c): endothelial cells(PECAM1, VWF), epithelial cells(EPCAM), fibroblasts(COL1A1, DCN), keratinocytes(KRT1, KRT14), lymphocytes(CD3D, CD3E), mast cells(TPSAB1), melanocytes(MITF, MLANA), monocytes(CD68, CD163, CD14), proliferative cells(PCNA, MKI67) and smooth muscle cells(ACTA2). In order to analyze the malignancy of each cell type, we further analyzed the copy number variation of each cell type using the mean expression of all cells as a reference. Proliferative cells derived from tumor samples have significantly higher copy numbers than other cell types and thus are identified as cancer cells (Fig. 4b).

**Fig. 4.**
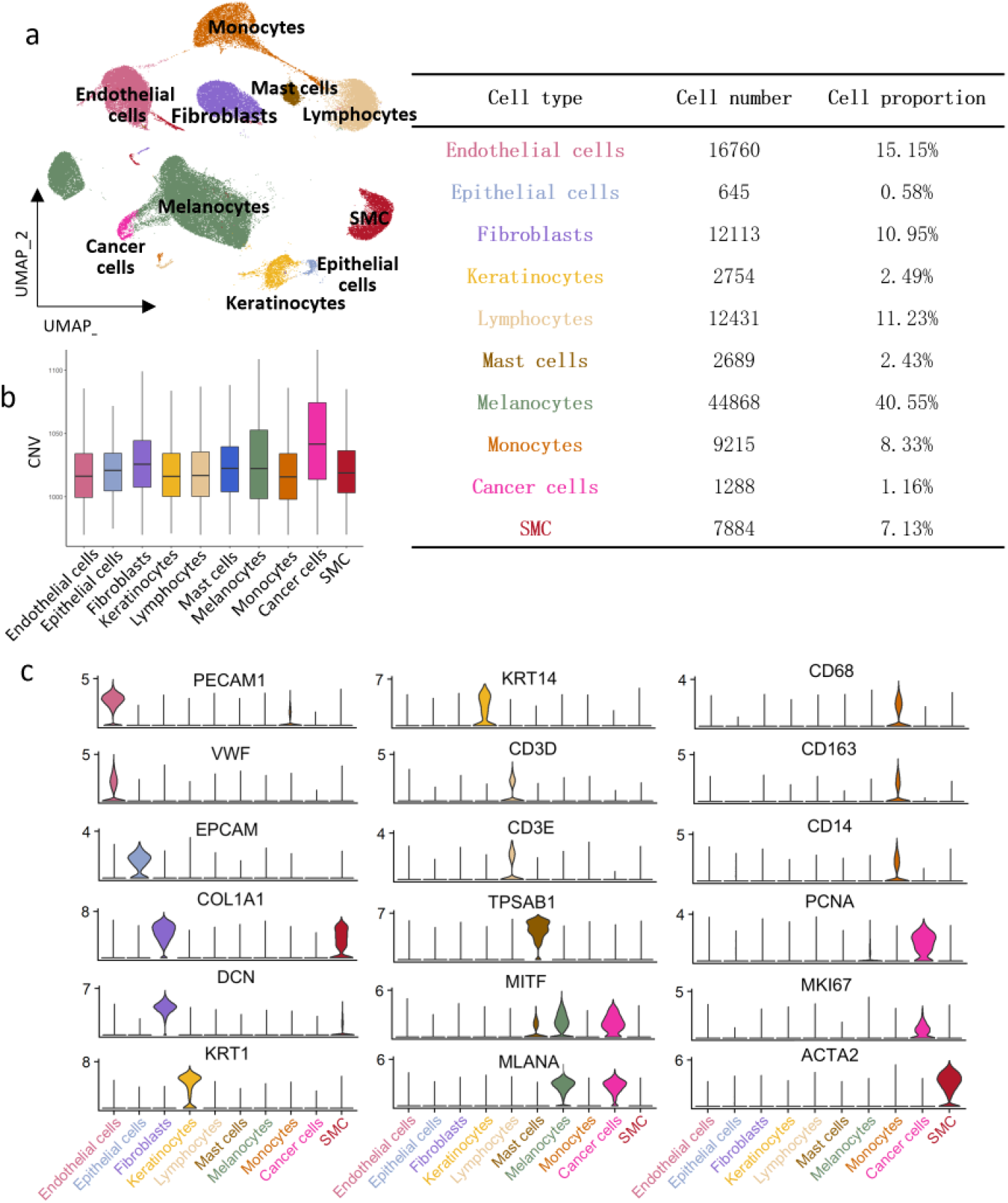
Single-cell atlas of acral melanoma. **a** Single-cell atlas of acral melanoma. **B** Copy number variations of each cell types. **c** Expression of canonical markers.

In section 2.2, we have discovered that subtype I is enriched with keratinocytes characteristics and has lower Breslow. It indicates that subtype I is less prone to deep invasion and thus tumor is confined to the epidermis, leading more keratinocytes from the epidermis to be easily mixed in clinical samples. However, with the limitations of traditional RNA-seq sequencing, only mixed values of various cells are obtained, which fails to accurately reflect gene expression at the cellular level. Hence, analysis based only on bulk data is susceptible to cellular composition. To test this hypothesis, we apply single-cell data and analyze the expression distribution of upregulated genes in subtype I obtained from bulk data. It is evident that the significantly upregulated genes of subtype I comprise a large number of keratinocyte-specific markers (Fig. 5a, b). However, keratinocytes cannot truly reflect the internal mechanism of tumorigenesis, and the difference in the proportion of keratinocytes is only a concomitant phenomenon. It is necessary to incorporate cell-level data to reveal the essential difference between the two subtypes. However, due to the high cost of single-cell sequencing omics data, the sample number is always limited. To solve this problem, we combine bulk data and single-cell data to remove the influence of noising features while achieving a balance of data precision and sample sizes.

**Fig. 5.**
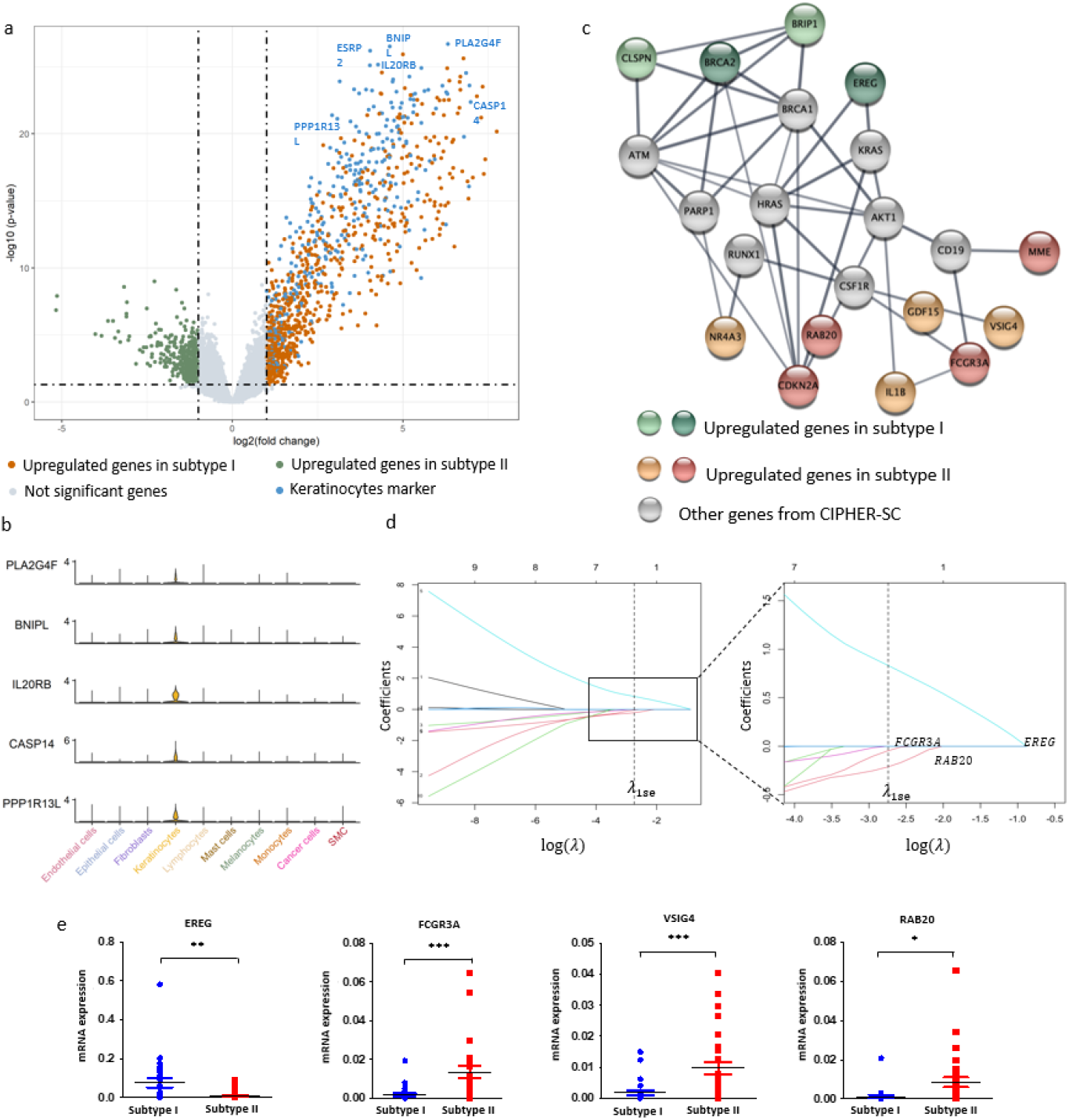
Identification of subtype markers. **a** Differentially expressed genes of two subtypes. Orange dots indicates the genes overexpressed in subtype I, green dots are the genes upregulated in subtype II and grey dots are genes not significantly differentially expressed from RNA-seq analysis. Blue dots are markers of keratinocytes calculated by single-cell RNA-seq analysis. **b** Expression distribution of keratinocytes markers. **c** Network related with subtyping of acral melanoma. **d** The lasso regression analysis for searching the optimal marker panel to distinguish the two subtypes. **e** RT-qPCR results to validate the expression of marker panels in clinical samples.

#### 2.4.2 Network-based disease gene prediction

In order to systematically analyze the relationship between diseases and genes, and improve the generalization of omics data analysis[24, 25], we also introduce network-based disease gene relationship inference algorithms CIPHER[26] and CIPHER-SC[27]. CIPHER algorithm reveals a hierarchical modular relationship between leimacro-phenotypes and micro-molecules in biological networks and can quantitatively describe the global disease-gene relationships. CIPHER-SC is an extended version based on CIPHER, which not only introduces cellular information, but also achieves high-precision cell-specific desease-gene assocation prediction through graph convolution on a context-aware network. These CIPHER series algorithms break through the bottleneck of multi-scale information fusion in complex systems and solves the problem of “small sample learning” widely existing in biomedical data, and achieve the most accurate disease-causing gene prediction.

#### 2.4.3 Subtype marker identification combining network-based prediction and omics analysis

Firstly, we analyze RNA-seq data to obtain the upregulated genes of two subtypes respectively (Fig. 3a). Secondly, highly expressed genes in tumors are selected by conducting differential expression analysis between acral melanoma and normal samples. Thirdly, to make up for the limitation of bulk RNA-seq data, we introduce single-cell transcriptome data to filter out keratinocyte markers for subsequent Identification of markers truly correlated with tumorigenesis. It’s worth noting that the clustering results do not change after removing the impact of keratinocyte markers. Fourthly, the network-based disease gene prioritization methods CIPHER and CIPHER-SC are applied and the top 10% predicted results are considered high-risk genes. In view of the above four points, subtype-related network is constructed from which 12 candidates markers are finally obtained: CDKN2A, BRCA2, FCGR3A, BRIP1, NR4A3, GDF15, CLSPN, IL1B, RAB20, MME, EREG, VSIG4 (Fig. 5c).

In order to further screen out the final subtype markers, we adopts the Lasso regression method to reduce structural risk while minimizing empirical risk (Fig. 5d). Five-fold cross validation is conducted and mean square error(MSE) is calculated under different value of *λ. λ*_min_ can be obtained when MSE reaches the minimum value. To avoid overfitting, we choose *λ*_1*se*_ that is one standard deviation away from *λ*_min_ and obtain 3 genes: EREG, RAB20 and FCGR3A. Among them, the coefficient of EREG is positive indicating it is a marker of subtype I, while on the contrary, RAB20 and FCGR3A are markers of subtype II. Since subtype II is of high risk, we add VSIG4 that highly expressed in subtype II for better identification. The marker panel consisting of 4 genes reaches as high as 0.946 of AUC in distinguishing the two subtypes. RT-qPCR is later applied to validate that EREG is overexpressed in subtype I while RAB20, FCGR3A and VSIG4 are overexpressed in subtype II (Fig. 5e).

### 2.5 Subytpe markers are specifically expressed in SPP1+ macrophages and FCN1+ macrophages respectively

Based on the acral melanoma single-cell atlas, we analyze the expression of subtype markers at the cellular level. It can be found that all markers are specifically highly expressed in monocytes (Fig. 6a). Therefore, we first extract the monocytes for further analysis. According to canonical markers, monocytes can be divided into dendritic cells(CD207, LILRA4, CD1C, LAMP3) and macrophages(CD68, CD163, CD206). Macrophages can be further subdivided into SPP1+ macrophages and FCN1+ macrophages [28]. It is evident that subtype I marker EREG is specifically expressed in FCN1+ macrophages, while subtype II markers FCGR3A and VSIG4 are specifically expressed in SPP1+macrophages (Fig. 6b). Based on the expression of markers of acral melanoma subtypes and macrophage subgroups, we applied correlation analysis and found distinct modularity. The correlation between EREG and markers of FCN1+ macrophages is higher, while VSIG4 and FCGR3A are highly correlated with markers of SPP1+ macrophages, indicating the great importance of macrophage subgroups in acral melanoma subtyping (Fig. 6c).

**Fig. 6.**
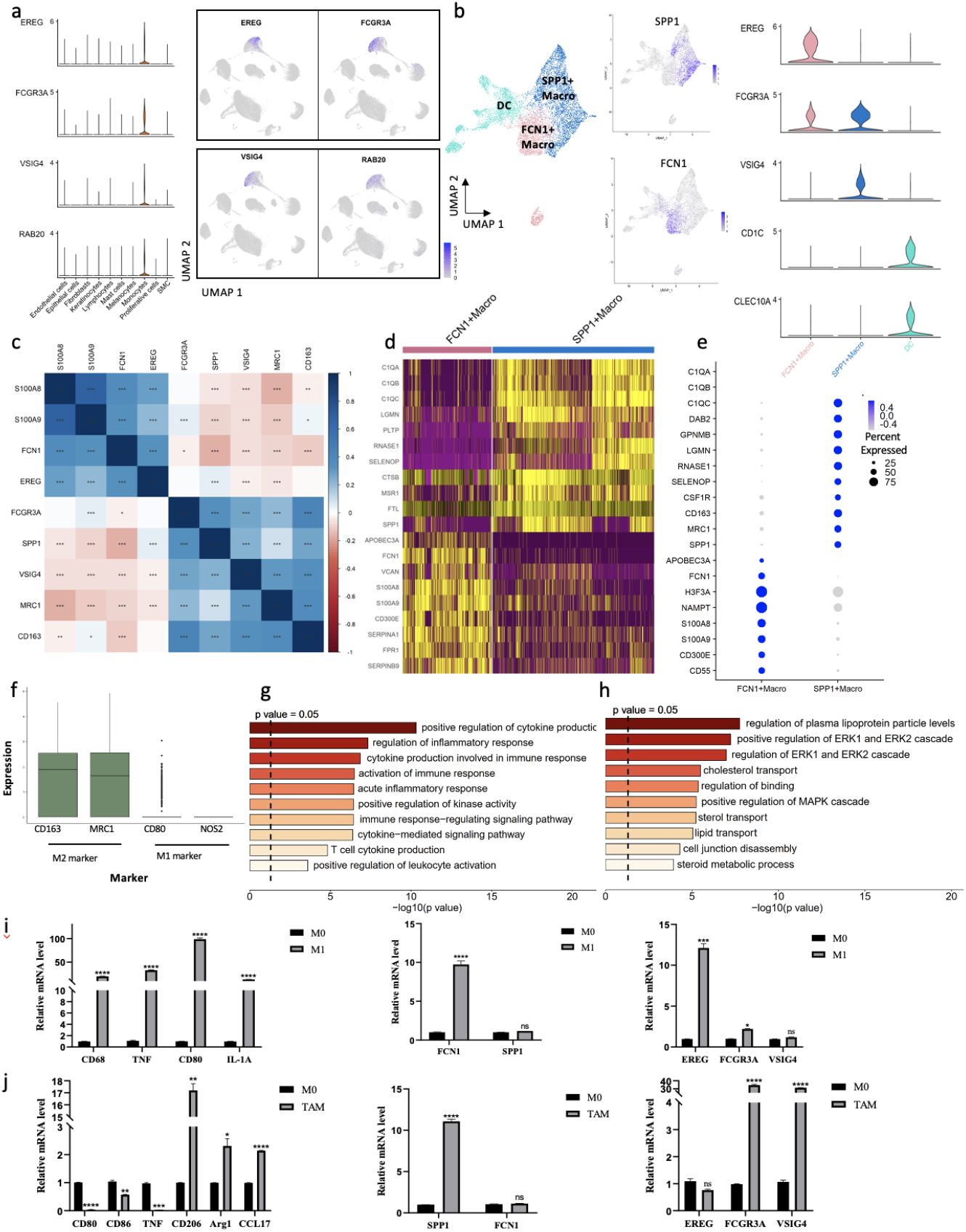
The function of two macrophage subgroups. **a** Expression of subtyping markers in single-cell RNA-seq data. All markers are specifically highly expressed in monocytes. **b** Subgroups of monocytes. Monocytes are further divided into DCs, FCN1+ macrophages and SPP1+ macrophages according to canonical markers. Subtyping markers are upregulated in FCN1+ macrophages and SPP1+ macrophages respectively. **c** Pearson correlation coefficient matrix of subtyping markers and macrophages marker genes. The shade of color indicates the strength of the correlation. Blue indicates positive correlation and red indicates negative correlation. **d**,**e** Differentially expressed genes of FCN1+ macrophages and SPP1+ macrophages. **f** Expression in SPP1+ macrophages of canonical markers of M1 macrophages and M2 macrophages. **g, h** Enriched GO functions of FCN1+macrophages and SPP1+ macrophages respectively. **i, j** qRT-PCR analyses of relative gene mRNA level in THP-1 cells, including M1 or M2 macrophages markers (CD68, CD80, CD86, TNF, IL-1A, CD206, Arg1, CCL17, FCN1 and SPP1) and subtyping markers (EREG, FCGR3A and VSIG4). Data are presented as mean *±* SEM from three independent experiments.

To compare the functions of these two macrophage subgroups in the development of acral melanoma, we calculate the feature genes of these two macrophage subgroups based on differential expression analysis and obtain their GO functions based on enrichment analysis. It is illustrated that FCN1+ macrophages highly express S100A8, S100A9, CD300E and etc (Fig. 6d,e). Their functions are mainly enriched in the immune-inflammatory response pathway, such as positive regulation of cytokine production, regulation of inflammatory response, cytokine production involved in immune response and etc (Fig. 6g). They are associated with immune activation in the tumor microenvironment and tends to function as classically activated macrophages(M1 macrophages). On the contrary, SPP1+ macrophages significantly express tumor-associated macrophages markers: C1QA, C1QB, C1QC, CD163, MRC1(CD206) and etc((*FDR <* 1*e* - 100, |*log*_2_*FC*| *>* 1.2) (Fig. 6d,e). Among them, CD163 and MRC1 are markers of alternatively activated macrophages(M2 macrophages), while M1 macrophages highly expressed CD80, iNOS(NOS2) and etc. Evidently, SPP1+ macrophages overexpressed M2 macrophage markers but not M1 macrophage markers, suggesting SPP1+ macrophages tend to play the role of M2 macrophages (Fig. 6f). As for the functions, the enriched pathways of SPP1+ macrophages are correlated with cell proliferation, lipid metabolism, and metabolic regulation of the tumor microenvironment, such as positive regulation of ERK1 and ERK2 cascade, lipid transport, steroid metabolic process and etc (Fig. 6h). In conclusion, these two macrophage subgroups play the role of M1 and M2 macrophages respectively. It indicates that the markers we identified to discriminate two subtypes of acral melanoma express in two macrophage subgroups with distinct functions. Specifically, markers of low-risk subtype are enriched in macrophages that promote immune activation, while markers of high-risk subtype are enriched in macrophages that enhance metabolism and proliferation. The expression profile of subtype markers reveals that the mechanism difference of the two subtypes lies in macrophage subgroups with distinct function in the microenvironment.

We next aimed to validate the relevance of the acral melanoma subtype marker to M1 or M2 macrophages at the cellular level. THP-1 was polarized into M1 macrophages by incubation with IFN-*γ* and LPS for 48h. qRT-PCR showed that M1 macrophage-associated markers (CD80, TNF and IL-1A) were significantly upregulated. And FCN1 was also highly expressed in M1 macrophages. Further, subtype I marker EREG was highly expressed in M1 macrophages, while subtype II markers VSIG4 and FCGR3A were not affected (Fig. 6i). Meanwhile, for tumor association macrophages polarization, we cultured THP-1 cell using SK-MEL-28 conditioned medium for 5 days. qRT-PCR showed that M2 macrophage-associated markers (CD206, Arg1 and CCL17) were distinctly increased, while M1 macrophage-associated markers clearly decreased. Also, SPP1 was increased in TAMs. Moreover, subtype II markers VSIG4 and FCGR3A were highly expressed in TAMs, and the expression of subtype I marker EREG had no variation(Fig. 6j).

### 2.6 SPP1+ macrophages and cancer cells interactions may contribute to melanoma progression

It has been found that macrophage subgroups with distinct functions in the tumor immune microenvironment can guide the identification of acral melanoma subtypes. To figure out the impact of these two macrophage subgroups on tumor cells, we use CellChat to infer intercellular communications. Based on the ligand-receptor interactions databases, communication probability is calculated and interactions with statistically significance are picked out. The number and the strength of interactions between all cell types are shown in fig, respectively (Supplementary Fig. 1a, b). Tumor immune microenvironment plays an important role in cell communication, among which macrophages have interactions with all other cell types. To analyze the function difference of two macrophage subgroups correlated with two acral melanoma subtypes, the interaction patterns between two macrophage subgroups and other cell types are compared in detail. SPP1+ macrophages exert influence on tumor cells from various signaling pathways, such as TNF signaling pathway, SPP1 signaling pathway and CXCL signaling pathway (Fig. 7a). Among them, SPP1 signaling pathway has the strongest communicaiton probability. By expressing ligand SPP1, SPP1+ macrophages communicate with other cells(tumor cells, melanocytes, epithelial cells, mast cells, etc) that express the corresponding receptors(CD44, ITGAV, ITGB5) (Fig. 7a, Supplementary Fig. 1c). It is worth noting that SPP1+ macrophages have the greatest impact on tumor cells, while FCN1+ macrophages have weaker interactions with tumor cells (Fig. 7b). To be specific, SPP1 and CD44 expressed by SPP1+ macrophages and tumor cells are the ligand-receptor interaction pair that obtain the highest score. Besides, SPP1-ITGAV, SPP1-ITGB1, and GRN-SORT1 also contribute to the communication between SPP1+ macrophages and tumor cells. It implies that SPP1 play an important role in the communicaiton with tumor cells.

**Fig. 7.**
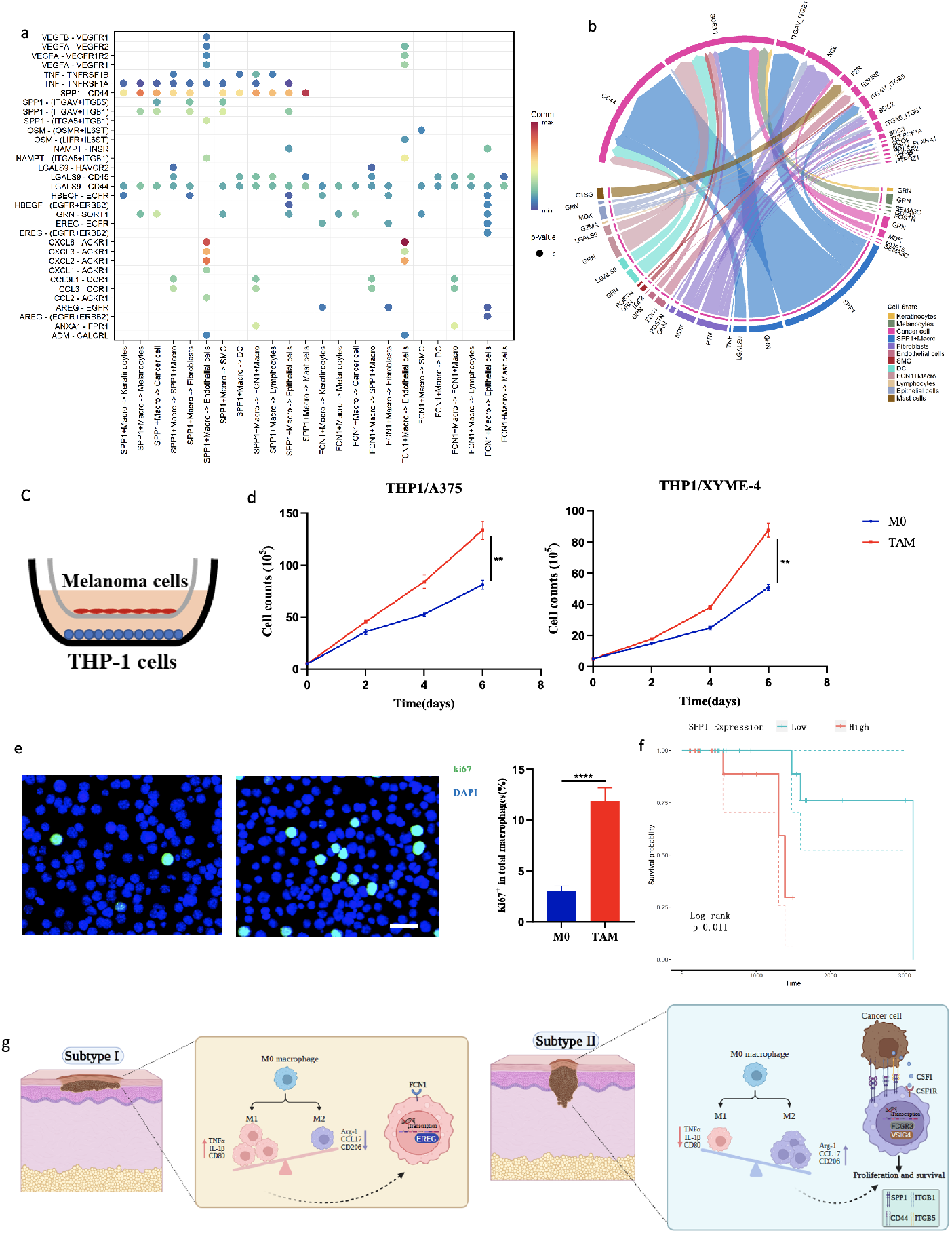
Intercellular communicaiton of two macrophage subgroups. **a** The cell-cell communication mediated by multiple ligand-receptor pairs from two macrophage subgroups to other cell groups. The color depicts communication probabilities and the dots indicates significant interactions. **b** Chord diagram to visualize the impact of all cell groups on cancer cells. SPP1+ macrophages exert the largest impact on cancer cells. **c** Schematic diagram of transwell co-culture experiments between THP-1 cells and melanoma cells (A375 and XYME-4 cells). THP-1 cells were seeded into the bottom chamber, and A375 or XYME-4 cells in the top chamber. **d** The number of THP-1 cells co-incubated with A375 (left) or XYME-4 (right) for different times. **e** Representative images of immunofluorescence staining (left) of ki67 in THP-1 cells co-cultured with or without XYME-4 cells for 3 days, and quantification of ki67+THP-1 cells (right). **f** KaplanMeier survival analysis of acral melanoma patients stratified by the expression of SPP1. **g** Action mechanism of different types of macrophages in two acral melanoma subtypes.

In addition to interacting with tumor cells, SPP1+ macrophages also communicate with mast cells and endothelial cells and play a vital role in various signaling pathways. We adopt the centrality metrics from graph theory to identify the dominant senders, receivers, mediators, and influencers for intercellular communications. As shown in (Supplementary Fig. 1d), mast cells are the dominant senders of CSF signaling pathways, while SPP1+ macrophages expressing the corresponding receptors CSF1R are the major receivers. CSF1 can recruit macrophages and promotes them polarized towards the M2 phenotype through interaction with its receptor CSF1R [29]. The results also account for the high expression of M2 macrophage markers in SPP1+ macrophages. In order to make the result conclusive, we verified the expression of CSF1R in M2 macrophages at the cellular level. We used SK-MEL-28 conditioned media to stimulate THP-1 cells for 5 days, which was induced into TAMs. And the result showed that CSF1R was significantly increased in TAMs (Supplementary Fig. 1e). SPP1+ macrophages interact with endothelial cells through the CXCL signaling pathway and act as the main senders. SPP1+ macrophages expressing ligands including CXCL8, CXCL3, and CXCL2 bind with the receptor ACKR1 on endothelial cells, which in turn promotes angiogenesis, tumor growth, and metastasis.

As shown, CSF1/CSF1R pathway could facilitate the proliferation of TAMs [30], and we showed that the expression of CSF1R is elevated in TAMs. Therefore, we wondered whether tumor cells could promote the proliferation of TAMs in acral melanoma. We selected two melanoma cell lines A375 and XYME-4, which is a primary acral melanoma cell established by our team previously [31]. We examined the effect of melanoma cells on macrophage proliferation using transwell co-culture system, in which macrophages were seeded into the bottom chamber with or without tumor cells into the top chamber. Melanoma cells promoted TAMs proliferation, either A375 or XYME in a time-dependent manner (Fig. 7c, d). Meanwhile, we collected macrophages co-cultured for three days for immunofluorescence staining. The result showed that proliferating macrophage (ki67+ cells) significantly increased in co-culture with XYME-4 (Fig. 7e). In addition, we explored the impact of TAMs, SPP1+ macrophages on acral melanoma. From a clinical point of view, high expression of SPP1 is also significantly correlated with poor patient prognosis (*P* < 0.02, logranktest) (Fig. 7f).

In conclusion, based on the single-cell data analysis of acral melanoma, it is found that the low-risk subtype I marker EREG is enriched in FCN1+ macrophages with the function of immune activation, while the high-risk subtype II markers FCGR3A and VSIG4 are enriched in SPP1+ macrophages associated with metabolism regulation. With the analysis of intercellular communicaiton, it is found that SPP1/CD44 signaling mediated by SPP1+ macrophages is vital for the tumor progression. The upregulated expression of SPP1 is significantly correlated with poor survival. Our findings reveals the internal mechanisms of subtype II acral melanoma and indicates that SPP1 is a potential therapeutic target for the FCGR3A+VSIG4+ acral melanoma subtype.

## 3 Methods

### Patients and sample collection

In this study, we totally collected tumor tissues and clinical information from 86 acral melanoma patients with informed written consent, and under approval of local medical ethnics. In the former RNA-seq and single-cell sequencing, there are 69 acral melanoma patients from multiple centers: 44 from Xiangya Hospital, 15 from Nanjing Skin Research Institute, and 6 from Hunan Provincial Tumor Hospital, and 4 from the Third Affiliated Hospital of Sun Yat-sen University. In the later validation, 17 acral melanoma patients were all from Xiangya Hospital. Fresh tissues were preserved in MACS tissue storage solution on ice and ready for transport.

### RNA-seq pipeline

Total RNA was extracted with MagZol reagent (Magen). After complete quality measurement, 1 *µ*g total RNA was used to enrich the mRNA with polyA tails by Oligo(dT) magnetic beads. In the presence of M-MuLV reverse transcriptase system, mRNA was reverse transcribed into cDNA, which was screened for 370 420 bp fragment and then amplified and purified. The library was constructed and sequenced using Illumina NovaSeq 6000 platform with 150 bp paired-end reads. FASTQ files were generated by CASAVA. To ensure the quality and reliability of the data analysis, the raw data needed to be filtered. It mainly included removing the reads with adapter, removing the reads containing N (N means that the base information cannot be determined), and removing the low-quality reads (the reads with Qphred ¡= 20 bases accounting for more than 50% of the whole read length). Meanwhile, Q20, Q30 and GC contents were calculated for clean data. Subsequent analyses were performed based on clean data with high quality.

### RNA-seq data analysis

We use functions implemented in edgeR R package to process the RNA-seq count data. Firstly, lowly expressed genes are filtered out by filterByExpr function and only protein-coding genes are kept for further analysis. Then, TMM(trimmed mean of M-values) normalization is applied to eliminate the biases betweeen libraries. Later on, QL F-test is performed to test the significantly differentially expressed genes(DE genes) between two clusters. To control the false discovery rata(FDR), we perform the Benjamini-Hochberg method and select genes with fold-change significantly above 1.2 at an FDR cut-off of 5%. After we obtain the DEGs, heatmap can be plotted using heatmap.2 function implemented in gplots R package. To analyze the function enriched in two clusters, we apply enrichGO function implemented in clusterProfiler R package which performs over-representation tests.

### Unsupervised clustering

In order to reduce data noise while reducing the computational load for clustering, we first select the top 500 genes according to the variance of gene expression in all samples, which can represent the intrinsic feature and difference between samples. Since no labels are provided in our data, we use K-Means algorithm to perform unsupervised clustering. Given a sample set *S* = {*S*_1_, *S*_2_, *s*_3_,, *S*_*Nm*_}, K categories are pre-defined. Then, by minimizing the within-class distance(equation 1), each sample is assigned a label.

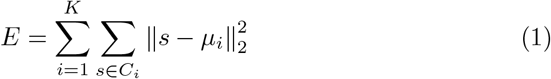

*µ*_*i*_ is the cluster centroids of the *i*th cluster. The optimal solution can be approximated by an iterative optimization. Firstly, K samples are randomly selected as the initial cluster centroids, and then the Euclidean distance between each sample and the K cluster centroids is calculated. the samples are classified into the cluster where the closest centroid is located. After that, we calculate the mean of each cluster as the new centroid. Finally, repeat the above steps until the coordinates of the K centroids converge. PCA is performed and the top two dimensions are selected for visualization.

To further determine the optimal number of the clusters, we apply Gap statistic and Silhouette Coefficient. The gap statistic method is based on hypothesis testing to compare the within-cluster sum of squares with its expectation under an appropriate null reference distribution. Silhouette Coefficient evaluate the classification effect based on the within-cluster distance and between-cluster distance. Suppose the average Euclidean distance between the sample *S*_*i*_ and samples of the same cluster is *a*_*i*_, and the average Euclidean distance between the sample *S*_*i*_ and samples in other clusters is *b*_*i*_. Silhouette Coefficient of *S*_*i*_ can be defined as:

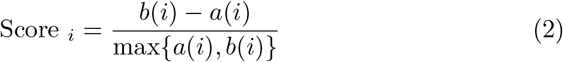

Then the final Silhouette Coefficient of the clustering can be calculated as the mean of Silhouette Coefficient of all samples 1/*N*_*m*_ ∑*Score*_*i*_. The optimal number of clusters is obtained when the silhouette coefficient reaches the maximum.

### Inference of cell type proportion

Tissues in different states or at different sites vary widely in the proportions of cell types. The cell-type composition of tissues has been found to be correlated with disease states and treatment responses. Thus, we employed Xcell to infer the proportions of cell types from bulk transcriptomic profiles and characterize the variation in the cellular composition between subjects. Xcell integrates expression profiles from six datasets including 64 immune and stroma cell types. By comparing to all other cell types, overexpressing genes in a certain cell type are identified, among which the most reliable signatures are selected later. We employed the single-sample GSEA to score samples and test the performance to select the most reliable signatures. Raw enrichment score of each cell type is defined as the average ssGSEA score calculated from the corresponding signatures of the cell type. After transforming to a linear scale and appling a spillover compensation, we can obtain the final enrichment score.

### Single-cell RNA sequencing pipeline

The acral melanoma tissues were extracted following surgical operation and washed in Hanks Balanced Salt Solution (HBSS) for three times. Then, single-cell suspensions were obtained by using The Human Tumor Dissociation Kit (Miltenyi Biotec) and gentleMACS Dissociator (Miltenyi Biotec). In addition, Dead Cell Removal Kit (Miltenyi Biotec, 130-090-101) was used to improve cell viability for single cell sequencing.

BD Rhapsody system was used for single-Cell RNA sequencing and library preparation in our study. Single cells from different patients were labeled with sample tags from the Human Immune Single-Cell Multiplexing Kit (BD Biosciences) based on the instructions. After washing with BD Sample Buffer and cell lysis, the cell capture beads were picked up for reverse transcription according to the manufacturer’s protocol (BD Biosciences, Single-Cell Capture and cDNA Synthesis). Single cell capture and cDNA library preparation were carried out following the BD Rhapsody Express Single Cell Analysis System (BD Biosciences). The libraries were loaded on an S1 flow cell (2100 cycle) and paired-end sequenced at *>*200,000 reads per cell depth on a Novaseq 6000 Sequencer (Illumina). PhiX (20%) was added to the sequencing run to compensate for the low complexity library[32].

### Single-cell sequencing data preprocessing

With the CWL-runner package, FASTQ files were processed according to the BD Rhapsody Analysis pipeline (BD Biosciences). Firstly, delete the low-quality reading pairs and analysis the R1 reads with quality filtering to recognize cell tags and UMI sequences. Then, the filtered R2 reads were mirrored to the transcriptome by using the STAR package. Reads with the same cell label, UMI sequence, and gene are wrapped into a single original molecule. Acquired events were adjusted by recursive substitution error correction (RSEC), an error correction algorithm (BD Biosciences) to rectify sequencing and PCR errors. The source of the sample is derived by using barcoded oligomeric antibodies (single cell multiplex kit; BD Biosciences).

We then use the standard pre-processing workflow encompassed in Seurat R package. We first visualize QC metrics and filter low-quality cells, empty droplets, cell doublets, and low-quality/dying cells with extensive mitochondrial contamination. After that, the NormalizeData method is applied to normalize the data. The cell expression profile is normalized by the total expression and then multiplied by a scale factor followed by a log-transforms operation. Next, we apply ScaleData function to shift the expression of each gene to a standard normal distribution so that highly-expressed genes do not dominate in the downstream analysis(such as dimension reduction and clustering). Later on, Principal Component Analysis is used to project the scaled data into the low-dimensional space and the top 30 principal components with the highest variance are remained. In the PCA space, Euclidean distance between cells is calculated to construct K nearest neighborhood network in which the weight is refined by Jaccard similarity. After that, FindClusters function implemented in Seurat is applied to cluster the cells. UMAP, a non-linear dimensional reduction technique is used for visualization.

### CNV analysis

InferCNV is applied to identify the evidence for copy number variations from single cell RNA-Seq data. In comparison to the average, CNVs for each region are estimated with the “denoise” settings and a value of 0.1 for “cutoff”. After that, the result is scaled to the range of −1 to 1 for each sample. The CNV score of each cell was calculated as quadratic sum to group deletions and duplications together.

### Lasso regression analysis

To select markers that best distinguish different clusters, we use Lasso regression analysis and employ glmnet R package which fits a generalized linear model (GLM) by penalized maximum likelihood 3. According to Occam’s razor principle, the structural risk is reduced on the basis of empirical risk minimization to enchance generalization of the model.

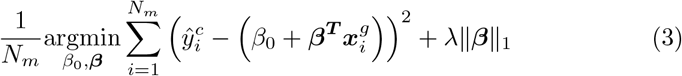

*N*_*m*_ is the sample size, 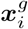 and 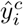 are gene expression vector and the cluster number of the *i*th sample. ***β*** is coefficient of the linear predictor part. *λ* is the regularization parameter that controls the overall strength of the penalty. To balance the trainning target and the structural complexity, we test the mean square error of the prediction at different value of *λ*.By conducting 5-fold cross validation,we can obtain the *λ*_*min*_ that minimizes the mean square error. To avoid overfitting, we finally use *λ*_1*se*_ one standard deviation from *λ*_*min*_.

### Intercellular communicaiton analysis

We use CellChat to quantitatively infer and analyze intercellular communication networks from single-cell RNA-sequencing data. To discover the cell-specific communications, we firstly identify overexpressed ligands and receptors in each cell type. Based on that, the overexpressed ligand-receptor interactions is obtained if either ligand or receptor is overexpressed. Then the biologically significant intercellular communication is inferred by assigning a probability value to each interaction and peforming a permutation test. The aggregated cell-cell communication network is obtained by counting the number of associations or adding the communication probability. The cell-cell communication at a signaling pathway level can be caculated by summarizing the communication probabilities of all ligands-receptors interactions correlated with the certain pathway.

### Cell lines and culture

XYME-4 cells were established by our team previously. Other cell lines were purchased from the American Type Culture Collection (ATCC). A375 and SK-MEL-28 were cultured in Dulbecco’s Modified Eagle Medium(BI), while THP-1 was incubated in Roswell Park Memorial Institute 1640(BI). All cell lines were supplemented with 10% fetal bovine serum (BI) and 1% antibiotics (penicillin/streptomycin) at 37°C in a 5% *CO*_2_ humidified incubator. THP-1 cell (8*105 per well) was seeded in 6-well plate and supplemented with PMA (100ng/mL) to adhere overnight at 37° in a 5% *CO*_2_ humidified incubator. For M1 polarization, THP-1 cell was cultured with recombinant human INF-*γ* (20 ng/mL, PeproTech) and lipopolysaccharide (100 ng/mL, Sigma) for 48h. For tumor associated macrophages (TAM) polarization, THP-1 cell was incubated with conditioned medium of SK-MEL-28 cell.

### qRT-PCR

Total RNA was extracted with TRIzol reagent (Bioteke Corporation), then retrotranscriped to cDNA using HiScript II Q RT SuperMix for qPCR (Vazyme) according to the manufacturer’s instructions. Then, 40 cycles of quantitative reverse-transcription PCR (qRT-PCR) were developed in 96-well plates using SYBR Green qPCR mixture (CWBIO) on the QuantStudio3 Real-Time PCR System. The fold change of gene expression was calculated by 2−(ΔCt _experimental group_ − ΔCt _control group_). All experiments were conducted three times independently. The sequence of primers was listed in Supplementary Table 1.

**Table 1.** Cox proportional hazards regression analysis of multiple cilnical characteristics

| Variables | Univariate analysis |  | Multivariate analysis |  |
| --- | --- | --- | --- | --- |
|  | Hazard ratio<br>95% confidence interval | P value | Hazard ratio<br>95% confidence interval | P value |
| Gender | 1.901<br>(0.209-17.270) | 0.568 | 0.630<br>(0.047-8.480) | 0.727 |
| Breslow | 0.630<br>(0.312-1.273) | 0.198 | 0.306<br>(0.088-1.057) | 0.061 |
| Clark | 1.198<br>(0.447-3.213) | 0.720 | 2.536<br>(0.542-11.871) | 0.237 |
| New subtype | 6.555<br>(1.031-41.660) | 0.046 | 27.134<br>(1.375-535.354) | 0.030 |

### Co-culture assay

THP-1 cells (2*105 per well) were seeded in 6-well plate and were induced to adhere overnight with PMA (100ng/mL). Then melanoma cells were seeded in the transwell-polycarbonate with 0.4*µ*m pores (Corning), and were co-cultured with THP-1 cells for different time (0, 2, 4 and 6 days). Finally, the number of THP-1 cells were obtained by observing and counting under the optical microscope.

### Immunofluorescence assay

After co-cultured with XYME-4 cells for 3 days, THP-1 cells in the bottom chamber were cleaned and fixed in 4% paraformaldehyde on ice for 15 minutes, and infiltrated with 0.5% Triton X-100 for 5 minutes. After blocking with 5% bovine serum albumin (BSA) for 1 hour, cells were incubated with ki67 monoclonal antibody (1:100, FITC, eBioscience) overnight at 4°C, and then stained with DAPI(1:10, Servicebio) to visualize nuclear DNA. The images were captured by confocal laser scanning microscope(TCS-SP8; Leica Microsystems) and analysed.

### Statistical analysis

qRT-PCR analyses were performed with GraphPad Prism8.0 software. Two-tailed Student’s unpaired t test was conducted for dual comparisons. The data represent the mean±SEM for at least three independent experiments.

## 4 Discussion

The traditional classification of melanoma is generally based on tumor location or pathological features. However, due to the high heterogeneity of the molecular mechanism for tumors, the therapeutic effects of patients with the same disease may vary considerably. Molecular subtyping contributes to the deep understanding of the mechanisms of different subtypes and provides the personalized treatment. Up till now, there is still a lack of in-depth analysis of heterogeneity in acral melanoma. We catch up with a new strategy that combines the bulk RNA-seq and single-cell RNA-seq expression profiles to discover novel subtypes of acral melanoma and identify their biomarkers.

We first apply the K-means method to classify samples into two clusters based on bulk data. With the analysis of clinical data, it is found that subtype II is a high-risk subtype that has thicker Breslow, greater infiltration of M2 macrophages, and poorer survival. Single-cell RNA-seq data is then used to compare the two clusters from cellular perspectives. It is found that feature genes that distinguish two subtypes primarily are highly expressed in keratinocytes, indicating that marker identification only based on bulk RNA-seq data will easily be influenced by noises and that biologically significant biomarkers will be masked. Therefore, based on that, we incorporate the single-cell RNA-seq data and the CIPHER-SC disease gene prediction method to filter out noisy genes. After that, Lasso regression analysis is employed to identify more generalized and high-precision marker panels which achieve the AUC as high as 0.946 in distinguishing different subtypes. The four markers are then validated by RT-qPCR.

At last, we discover that the subtyping markers are specifically overexpressed in different subgroups of macrophages. As the most predominant immune cells in the tumor microenvironment (TME)[33], macrophages are recruited from circulating monocytes and tissue resident macrophages to TME and are preferentially polarized into the M2 macrophage via multiple cytokines and chemokines to develop into TAMs[34]. It has been discovered that the number of M2 macrophages increases in thick(*>*1mm), ulcerated, and highly mitotically active acral melanomas[35]. In our study, we find that markers of low-risk subtype I(EREG) are highly expressed in FCN1+ macrophages, while markers of high-risk subtype II(FCGR3A, VSIG4) are highly expressed in SPP1+ macrophages. In addition, these two subgroups of macrophages are prone to play the role of M1 and M2 macrophages respectively. Based on the intercellular communication analysis, it is revealed that SPP1+ macrophages communicate with tumor cells mediated by the SPP1/CD44 signaling pathway and can promote tumor growth. Apart from that, the high expression of SPP1 is significantly correlated with poor survival of acral melanoma patients. These findings reveal the intrinsic mechanism of the high-risk acral melanoma subtype and provide a new therapeutic target for the FCGR3A+VSIG4+ acral melanoma subtype.

## 5 Acknowledgements

This study was supported by grants No.81874138, 82073020, 62061160369 and 81225025 from General Program, The National Natural Science Foundation of China, Hunan Natural Science Foundation for Distinguished Young Scholars (Grant No.2021JJ10073)

